# MONDE·T: A Database and Interactive Webserver for Non-Canonical Amino Acids (ncAAs) in the PDB

**DOI:** 10.64898/2025.12.21.695100

**Authors:** Alex Waldherr, Gesa L. Freimann, Andrei N. Lupas

**Affiliations:** Department for Protein Evolution, Max Planck Institute for Biology, Max-Planck-Ring 5, 72076, Tu bingen, Germany

**Keywords:** protein, cell biology, synthetic biology, Ramachandran, protein structure, protein design, non-canonical amino acids, non-standard amino acids, database, webserver, post-translational modifications, biochemistry

## Abstract

**Summary:** Non-canonical amino acids (ncAAs) are understood as amino acids that are not genetically encoded. They are of wide interest for protein design and applications in pharmaceutical and cell biological research, but their impact on protein structure has not been explored systematically. We therefore collected all ncAAs in the Protein Data Bank (PDB) into the MONDE·T database, amounting to **1**,**875** different chemical types in >10,000 entries. They are made accessible through a webserver that allows for data download, visualization of the structures in which they occur, plots of their backbone torsion angles compared to canonical residues, and exploration of their similarity to these. Analyses can focus on a single ncAA or a specific protein structure, illustrated here by example applications.

**Availability and implementation:** MONDE·T is hosted at the Max Planck Institute for Biology Tu bingen, accessible online at https://mondet.tuebingen.mpg.de. The database is available for download under the Creative Commons 4.0 License.

## Introduction

Non-canonical amino acids (ncAAs) extend the repertoire of the genetically encoded canonical amino acids, both in the backbone and in the side chains. In the backbone, D-amino acids introduce mirrored enantiomers of the standard set, while *β*-amino acids introduce an additional carbon atom into the backbone, allowing for more flexible designs. The backbone can be further elongated by additional carbon atoms to yield γ-, δ-, ε-etc. amino acids. The backbone can also be elongated by the introduction of other elements between the amino and carboxylic groups.

As side chain modifications, nature already produces a wide range, both reversible and irreversible. For example, the most widespread by far is the reversible phosphorylation of serine, threonine and tyrosine residues in cellular signal transduction. ^1.2^ Reversible modification of lysine residues in histone tails by acetylation and methylation also produces essential biological outcomes.^3^ Prominent irreversible modifications are for example the systematic conversion of prolines to hydroxyprolines in collagens^4^, the N- and O-linked glycosylation^5.6^ of side chains during the maturation of surface proteins and the attachment of lipid tails to terminal cysteine residues^7^ as membrane anchors. Beyond these, chemistry provides an almost endless list of synthetic modifications, for example the use of nucleotide bases as side chains to produce PNAs, peptide nucleic acids.^8^

Such changes of chemical identity can influence the protein structure landscape in profound ways. For an example of backbone-dependent structural effect, the global structure and function of the ribosomally produced toxin polytheonamide entirely depend on the systematic conversion of every second amino acid to the D-form by a processive epimerase. ^9^ Further extensive modification of the side chains allows the toxin to fold into a membrane spanning ion channel that provides the toxic function. ^10^ The same structure and toxicity can be brought about in gramicidin, which is produced by non-ribosomal peptide synthases. ^11^ For an example of side chain dependent structural effect, a new form of coiled coil not previously seen in nature could be generated by the systematic inclusion of the non-canonical side chains alloisoleucine and norvaline. ^12^ The expectation in the field of protein design is that modified amino acids open entirely new areas of protein structure, both in the static and dynamic behavior of protein folds. The extent to which this expectation can be realized hinges critically on an understanding of the effects of non-canonical amino acids on protein structure.

As a first step towards this goal, we set out to collect a complete set of all ncAAs in experimentally verified protein structures. Previous efforts, such as the iNClusive database^13^ (466 ncAAs), are incomplete and lack structural context. Unexpectedly, we found that a high count of ncAAs were already included in proteins and are structurally determined: **1**,**895** unique ncAAs (**94x the canonical diversity**) in 23,149 structures. We curated this dataset into MONDE·T, a database and webserver, with tools for interactive analysis. Users can explore 1D sequences, 2D Ramachandran landscapes, 3D structural models, and chemical similarities. Our focus on experimental data makes MONDE·T a useful tool for the application in biomedical research.

## Results

### Data Curation

For the MONDE·T dataset, we aimed to identify all cases in which ncAAs were incorporated into a polypeptide chain, but the PDB contains many other compounds in addition to polypeptide chains.

We therefore used the worldwide chemical component dictionary (https://www.wwpdb.org/data/ccd) as a starting point, filtered out any unbound or bound ligands and separated true backbone inclusions in peptide backbones from those found in other chain types, like nucleic acids.^14,15,16,17^ In quantitative terms, we filtered **48**,**611** unique chemical components to a set of **2**,**762** found in a polymer sequence, of which **1**,**895** are found within peptide backbones. Of these, 20 represent the canonical set, leaving **1**,**875** ncAAs including terminals (e.g. amides). For each ncAA, we extracted all chain occurrences, further annotating these with source organism, experimental method, resolution, and polypeptide chain sequence in single letter code (in which ncAAs are denoted by **X**). Additionally, we included a version of the sequence in which ncAAs are represented by their chemical component ID (their compound abbreviation in PDB). Note that these are usually three letters long but may occasionally contain five letters. Note further that an overview page for a given compound in PDB can be accessed under https://www.rcsb.org/ligand/XXX(XX).

The MONDE·T dataset is available under the Creative Commons 4.0 License and updated half-annually. Filtered subsets or the whole annotated dataset can be copied to the clipboard or downloaded as CSV files via the webserver.

### Webserver

The webserver offers four functional areas, accessible from tabs at the top of the webpage.

#### Ramachandran plot tab

The Ramachandran plot lies at the core of MONDE·T, because the backbone information provided by this plot is central to understanding the structural properties of a polypeptide chain and its residues. Every residue in a polypeptide chain can be described by the angle *ϕ* to its preceding neighbor and the angle *ψ* to its succeeding neighbor. In a Ramachandran plot, the *ϕ*-angle is shown on the *x*-axis and the *ψ*-angle on the *y*-axis. To indicate how the canonical amino acids shape protein backbones, selection of radio-buttons in the field “*Settings* → *Select canonical background*” provides *(ϕ, ψ)*-angles in PDB entries with a resolution ≤ 1.2 A and R_observed_ ≤ 0.2 (1,511,165 residues in total).

To analyze ncAAs against this background, users can enter a chemical component ID in the field “*Settings* → *Select non-canonical amino acid”* or provide a protein structure in the field “*Settings* → *Select PDB ID”*, either by uploading it as a PDB/MMCIF file or by fetching it via its PDB ID.

If a chemical component ID was entered, a scrollable table of all PDB entries containing this ncAA pops up. This table is interactive. By clicking on PDB entries in this table, the backbone angle for the ncAA in this entry is displayed as a red star on the Ramachandran plot to the right, against the background chosen by the user. To display the complete experimentally determined coverage for this ncAA, a slider can be dragged to select or deselect ranges of entries (be aware that terminal residues either lack a *ϕ-* or *ψ*-angle and are therefore not plotted in the Ramachandran space).

If a PDB ID is entered or a PDB/MMCIF file is uploaded, the chains of the structure will be listed below the Ramachandran plot. By selecting a chain from this list, the user can build the plot for a region of interest using a second slider. The plot is interactive. Canonical residues are displayed in blue, ncAAs in red. More labels are displayed upon hovering, listing residue code and number. The curated plot can be “saved to disk” as PNG.

Since the analysis can be performed irrespective of whether an uploaded structure contains ncAAs, this tool offers rapid and intuitive access to the backbone geometry of any polypeptide chain.

#### Database tab

This tab provides access to the MONDE·T dataset structured by chemical component ID. If no ID is entered, the table lists all chemical components in the database with their annotations in individual columns. The table can be reconfigured by sorting a specific column and can be filtered by a search term. If a component ID is entered, the table displays this entry with its annotations. The dataset, complete or filtered, can be downloaded as CSV file.

#### Chemical similarity tab

This tab compares ncAAs to the twenty canonical amino acids according to their Tanimoto similarity. A table is searchable by chemical component ID or other text strings and sortable by columns. Two canvases are present to visually compare 2D structures.^18^ When an ncAA was selected in the *Ramachandran plot tab*, its 2D chemical structure is pre-rendered into one of these.

#### Protein structure tab

Another way to obtain information about ncAAs in polypeptide chains is to interactively explore the 3D structure. The tab uses Mol* Viewer^19,20^ (as in PDB) to display the structure uploaded in the *Ramachandran plot tab* in gray colors, with non-canonical positions highlighted in red.

### Application Example: Posttranslational Modifications

A widespread natural modification is the phosphorylation of residue side chains. If we wish to track structural examples of histidine phosphorylation (i) we can query the *Database tab* with the search term *histidine* and (ii) obtain two IDs with a **‘-phosphono-’** name prefix: **HIP** (1-phosphonohistidine, 16 entries) and **NEP** (3-phosphohistidine, 68 entries). (iii) These we fetch as ncAAs in the *Ramachandran plot tab* and visualize their backbone torsion angles against a selected canonical histidine background. (iv) We observe (see Fig. 2 top row) that **NEP** spreads in the canonical histidine space, whereas **HIP** is concentrated at its periphery. (v) We can further compare the phosphorylated **HIP** to the histidine methylated at the same position, **MHS** (1-methylated histidine, 67 entries). We find that **MHS** occupies a Ramachandran space that differs substantially from the canonical one.

Posttranslational modifications can also be introduced synthetically, for example with the serine modifier [dimethylamino(ethoxy)phosphoryl]-formonitrile. This compound, notorious as the nerve agent **tabun**, acts by covalently modifying the catalytic serine in acetylcholinesterase. In MONDE·T, we find this modification for example by querying the *Database tab* with serine, ordering the resulting table by *total_counts*, and identifying the correct compound, **SUN** (9 entries). We observe that the modified serine occupies an unusual Ramachandran space at the edge of the canonical distribution (Fig. 2 center left), but that the equivalent unmodified serine (e.g. PDB ID **2HA3**, pos. 203) is found in the same space.

Workflow: *database prefiltering* → *concrete identification of IDs and annotation data of interest* → *utilization of ID in the plotting modality* → *comparison of related chemical variants*.

**Figure 1:**
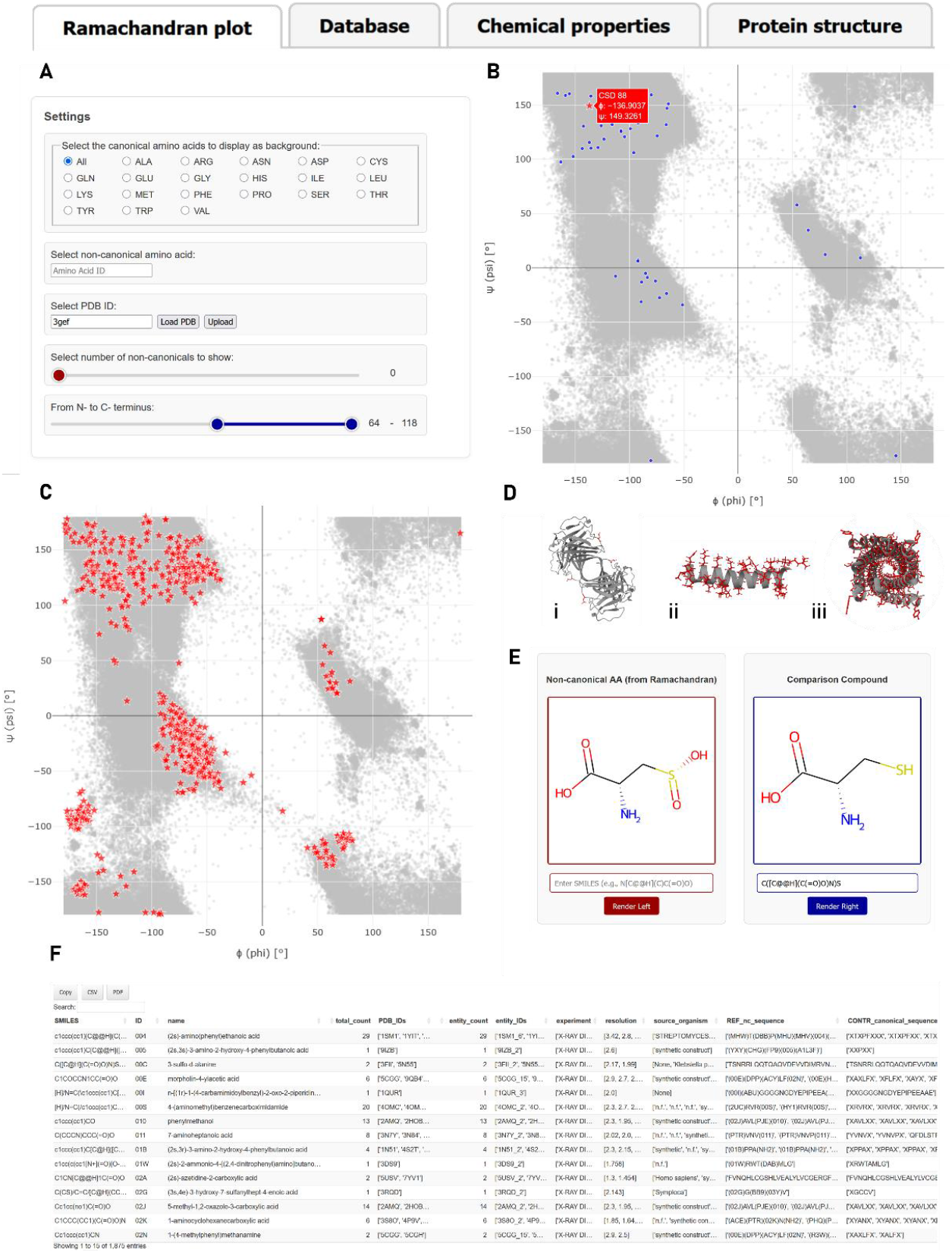
A collage of functionalities in MONDE·T. MONDE·T is split into four tabs: Analysis starts in the *Ramachandran plot tab*. **A:** A PDB/MMCIF file can be uploaded or fetched (makes plot **B**, exemplified with PDB ID 3GEF). Alternatively, ncAAs of interest can be fetched via their chemical component ID and their occurrences in PDB get plotted as landscape (makes plot **C**). **D:** If a protein was fetched, its structure is shown as 3D view in the *Protein Structure tab*. For fast analysis, the ncAA residue are highlighted as red sticks, exemplified here for (i) 3GEF (same structure as in B), (ii) the toxin polytheonamide B, (iii) the designed D-AIB-3_10_ coiled coil. **E:** If an ncAA is analyzed, this compound of interest is prerendered in the *Chemical Similarity tab*, which allows for visual comparison with another component. **F:** All data can be searched and downloaded in the *Database tab*.

**Figure 2:**
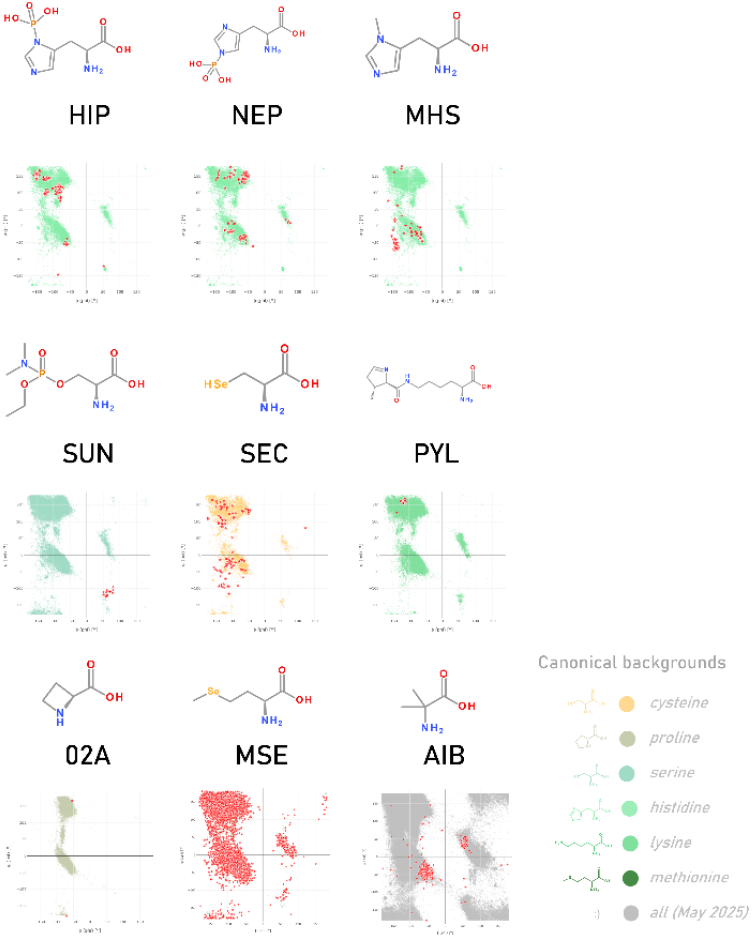
A collage of the ncAAs mentioned in the application examples in MONDE·T style. ncAAs are displayed as red stars over a canonical background of choice. This allows for comparison of whether the ncAA fold landscape overlaps with canonical torsion angle distributions.

### Application Example: Stop Codon Reassignment

Several ncAAs are introduced biologically by rededication of stop codons, for example selenocysteine^21^ (**SEC**, 93 entries) and pyrrolysine^22^ (**PYL**, 9 entries), through highly specialized machinery directly at the ribosome. In the *Ramachandran plot tab*, we select cysteine or lysine backgrounds and plot the chemical IDs **SEC** or **PYL** across all their PDB occurrences against these (Fig. 2 center right). We observe that **PYL** is fully contained within the canonical region, but **SEC** drifts into the non-canonical space already seen for **MHS** above.

The same strategy of codon reassignment has been adopted by researchers to incorporate ncAAs for practical purposes. Because of the effort to introduce additional tRNAs and orthogonal synthetases which can decode the suppressed stop codon, this method has been mostly used for functional goals and there are few structures in PDB. One of the few examples, where 2-cyanophenylalanine (**9IJ**, 16 entries) was included to probe the photosynthetic reaction center of *Rhodobacter sphaeroides* (PDB ID **8VTJ**), shows complete occupancy of the ncAA residues inside the canonical Ramachandran space as was verified by these authors to support their analysis on isolated functional effects. ^23^

Workflow: *known chemical component ID* → *optional: select respective canonical background* → *fetch by chemical ID* → *plot and check Ramachandran landscape of component*.

### Application Example: Amino Acid Analogs

Biological systems have evolved a number of amino acid analogs that are incorporated at the ribosome with toxic effects. For example, the toxin of lily-of-the-valley flower, (2*S*)-azetidine-2-carboxylic acid is a proline-analog lacking one carbon in the ring.^24^ It is misincorporated by the ribosome instead of proline and has teratogenic effects, severely affecting collagen structure.^25,26^ A search for (2*S*)-azetidine-2-carboxylic acid in the *Database tab* yields the abbreviation **02A** (2 entries) and its comparison with canonical proline in the *Ramachandran plot tab* shows **02A** at the edge of the proline distribution (Fig. 2 bottom left). However, uploading collagen structures for comparison (e.g. PDB ID **1K6F**) shows **02A** in the same space, suggesting that its toxic effects are not due to changes in the backbone torsion angles. However, there are no collagen structures incorporating **02A** in PDB at present.

Synthetically, the incorporation of amino acid analogs by mistranslation of sense codons is generally pursued with selective pressure incorporation, in which the organism is exposed to media with the ncAA replacing the canonical analog.^27^ The most widespread application of this is the incorporation of selenomethionine for crystallographic structure determination and correspondingly **MSE** is the most common ncAA in MONDE·T (10,214 entries). As fully expected, since **MSE** is used in place of methionine precisely because its structural properties are the same, a comparison in the *Ramachandran plot tab* shows identical distribution (Fig. 2 bottom center).

### Application Example: Chemical Peptide Synthesis

Evidently, chemical peptide synthesis is not a method inspired by natural processes. The incorporation of ncAAs by this method could be fundamentally separated into two areas, site-specific optimization of pharmaceutical candidates or global scaffold design for protein engineering. An example for site-specific optimization is the exchange of an alanine residue by α-aminoisobutyric acid (**AIB**, 201 entries) in the currently widespread GLP-1 agonist semaglutide, to prevent cleavage by dipeptidylprotease 4.^28,29^ Comparing a semaglutide structure (e.g. PDB ID **7KI0**) in the *Ramachandran plot tab* against the alanine background shows its **AIB** residue in the canonical space of α-helices.

Comparing the complete **AIB** landscape across all 201 occurrences in the *Ramachandran plot tab* indicates a strong preference of **AIB** overall for the L- and D-α-helical regions (Fig 2 bottom right).

Indeed, as a non-chiral amino acid, the Ramachandran space of **AIB** is equivalent in L- and D-configurations, making it very attractive for the design of new protein structures not accessible with the canonical amino acid set. Case in point, building coiled coils from 3_10_–helices, rather than α-helices, was only possible in D-configuration with a third of the residues being **AIB** (PDB ID: **7QDI**).^30^

## Conclusion

The MONDE·T toolkit is an entry point for the exploration of ncAAs within polypeptide chains. The MONDE·T database is built entirely on proteins of known structure that contain ncAAs. At the center of analyses performed through MONDE·T is the Ramachandran representation of backbone torsion angles, providing an instant 2D indicator of how amino acids shape protein backbones. This tool is usable for any polypeptide chain, whether or not it contains ncAAs, providing rapid and interactive access to the Ramachandran space in general. Access to MONDE·T is possible both through chemical component IDs and through PDB/MMCIF-format structures, which can either be uploaded by the user or fetched from PDB. We anticipate that MONDE·T will be useful to research scientists who are considering use of ncAAs for biomedical inquiry.

### Update commitment

We commit to updating the database semi-annually. As a rough growth estimate: Between April 2025 and October 2025, the wwPDB chemical component dictionary gained **1**,**936 entries** (+4%) of which MONDE·T gained **52** ncAAs and capping groups (+3%).

### Maintenance commitment

We commit to keeping the MONDE·T database openly available and updated at the Max Planck Institute for Biology for the next two years.

## Author contributions statement

AW compiled the database and code files for MONDE·T functionalities. AW and GLF implemented the backend of the webserver and designed the analysis functionalities. AW, GLF, ANL wrote and reviewed the manuscript.

## Acknowledgments

The authors thank Dek Woolfson for valuable improvement suggestions and the open-source communities of BioPython, RDkit, and Mol* Viewer for software development and Johannes Wo rner for reliable IT infrastructure and maintenance. This work was supported by institutional funds from the Max Planck Society. AW and GLF are members of the International Max Planck Research School ‘From Molecules to Organisms’.

